# Single-cell multi-omic characterization of the soybean root response to cyst nematode infection

**DOI:** 10.64898/2025.12.05.692575

**Authors:** Xuan Zhang, Vinavi A. Gamage, Xunliang Liu, Ziliang Luo, Mark A.A. Minow, Hao Zhang, Melissa G. Mitchum, Robert J. Schmitz

## Abstract

Soybean cyst nematodes parasitize soybean roots by inducing the formation of multinucleate syncytia to feed and complete their life cycle. However, the molecular basis of syncytia initiation and development remains limited. Here, we generated an integrated single-nucleus RNA and chromatin accessibility profile of soybean roots from infected and uninfected plants. We profiled 56,448 high-quality nuclei and identified all major root cell types, including distinct subpopulations enriched with syncytial nuclei. Transcriptomic and chromatin accessibility analyses support their procambium cell signature that progresses through distinct stages associated with immune suppression, cell fusion, and endoreduplication. Integrative gene expression and transcription factor motif chromatin accessibility analyses identified CAMTA1, as a potential transcriptional repressor of defense-related genes including receptor-like kinases, whereas MYB, MYB-related, and E2F were identified as potential transcriptional activators that coordinate cell wall remodeling, chromatin modification and DNA replication, respectively, across developmental trajectories. These findings provide mechanistic insights into host reprogramming during nematode parasitism and serve as a foundational resource for engineering nematode-resistant soybean.

## Introduction

Soybean (*Glycine max [L.] Merr*.) is a globally important legume crop, serving as a key source of plant-based protein and oil for human consumption and livestock products [1]. However, soybean yield losses exceed $1.5B annually due to the soybean cyst nematode (SCN, *Heterodera glycines*), making it the most economically damaging soybean pathogen [2,3]. SCN management relies on non-host crop rotation and cultivar resistance [4]. Currently, genetic resistance is limited to several *Resistance to Heterodera glycines* (*Rhg*) genes and combinations thereof [5–9]. This reliance on limited *Rhg* genes in commercial soybean cultivars has led to widespread virulence in SCN field populations [10]. Complex nematode virulence dynamics and an incomplete understanding of the molecular basis of soybean-SCN interactions have limited the pace of novel resistance improvements [11]. Thus, a deeper understanding of syncytia formation, specifically how host cells are reprogrammed during nematode infection, is necessary to enable durable resistance breeding.

A critical step in SCN parasitism is syncytia formation, specialized feeding structures derived from root cell reprogramming [12]. Upon penetration of the root by second-stage juvenile (J2) SCN, the nematode migrates toward the vascular cylinder and selects an initial feeding cell in the cortex, endodermis, or pericycle [13]. Partial cell wall dissolution leads to the fusion of hundreds of neighboring cells, producing a multinucleate syncytium [14,15]. During syncytium formation, cells undergo extensive cytological reprogramming, including nuclear enlargement, endoreduplication, increased organelle proliferation and metabolism, and remodeling of cell walls to form elaborate wall ingrowths to facilitate solute transport [16]. Despite advances in bulk transcriptomic and proteomic studies, the molecular mechanisms underlying syncytium development and epigenetic reprogramming are poorly characterized.

Bulk-tissue transcriptomics has examined the soybean-nematode interaction [17–20], but have had limited ability to measure syncytia gene expression. More recently, laser-capture microdissection (LCM) transcriptomic measurements successfully isolated syncytial cells, garnering insights into syncytial gene expression at different developmental stages [15,21–23]. However, these LCM studies lack whole-root context and inadequately capture the earliest stages of syncytia initiation. Single-cell technologies enable the study of plant-microbe interactions across all cell types in a tissue and can capture a developmental gradient from the initial point of infection [24]. For example, previous single-cell Assay for Transposase-Accessible Chromatin sequencing (scATAC-seq) and RNA-seq characterized the molecular signatures of infected soybean cells in developing nitrogen fixing nodules [25,26]. However, to date no single-cell multi-omic (i.e., transcriptome + chromatin accessibility) technology has been used to study plant-nematode interactions.

To better characterize soybean syncytium initiation and development, we applied single-nucleus RNA sequencing (snRNA-seq) and scATAC-seq to mock-inoculated and SCN-infected roots of Williams 82, a SCN-susceptible soybean cultivar. Profiling chromatin accessibility and gene expression in 56,448 individual nuclei, we characterized procambium-derived cell clusters undergoing distinct stages of syncytium development. Specifically, we identified three states: (i) early response to nematode infection, (ii) multinucleation via neighboring cell fusion and (iii) established syncytia undergoing endoreduplication. We further identified putative transcription factors underpinning the host cell reprogramming process: CAMTA1 transcriptionally represses defense-related genes, and MYB transcription factors activate genes involved in the cell-wall modification likely contributing to syncytium initiation and expansion. Additionally, we constructed a comprehensive syncytium developmental trajectory, revealing temporal changes in regulatory gene modules over the transition from early infected cells to mature syncytium. Collectively, this study provides novel insights into the molecular mechanisms underlying syncytial development and offers a valuable single-cell resolution resource for understanding plant-nematode interactions. These findings have practical implications for the genetic improvement of SCN resistance by pinpointing specific cell types and regulatory circuits as potential breeding or engineering targets.

## Results

### A single-cell multi-omic atlas of soybean roots during SCN infection

To dissect the molecular responses of soybean roots to SCN infection and track early syncytium development, we performed snRNA-seq and scATAC-seq on mock-inoculated and SCN-infected soybean roots from the susceptible genotype Williams 82 at 5 days post-inoculation (DPI) with two biological replicates (Fig. 1A, Methods). The soybean roots were inoculated with infective J2 and allowed to infect the roots for 24 hours after which the roots were washed to remove any J2 that had not yet penetrated the roots. This step allowed for the collection of root tissue with a higher percentage of sedentary life stages of SCN enabling the capture of a mix of initial syncytial cells through developing syncytia up to a maximum of ∼4 days old. After stringent quality control, we retained 33,441 high-quality scATAC-seq nuclei, with a median of 12,485 Tn5 insertions per nucleus, and 23,007 snRNA-seq nuclei, with a median of 1,271 UMIs and 1,038 detected genes per nucleus (Fig. S1A-H, Table S1,S2).

**Fig. 1.**
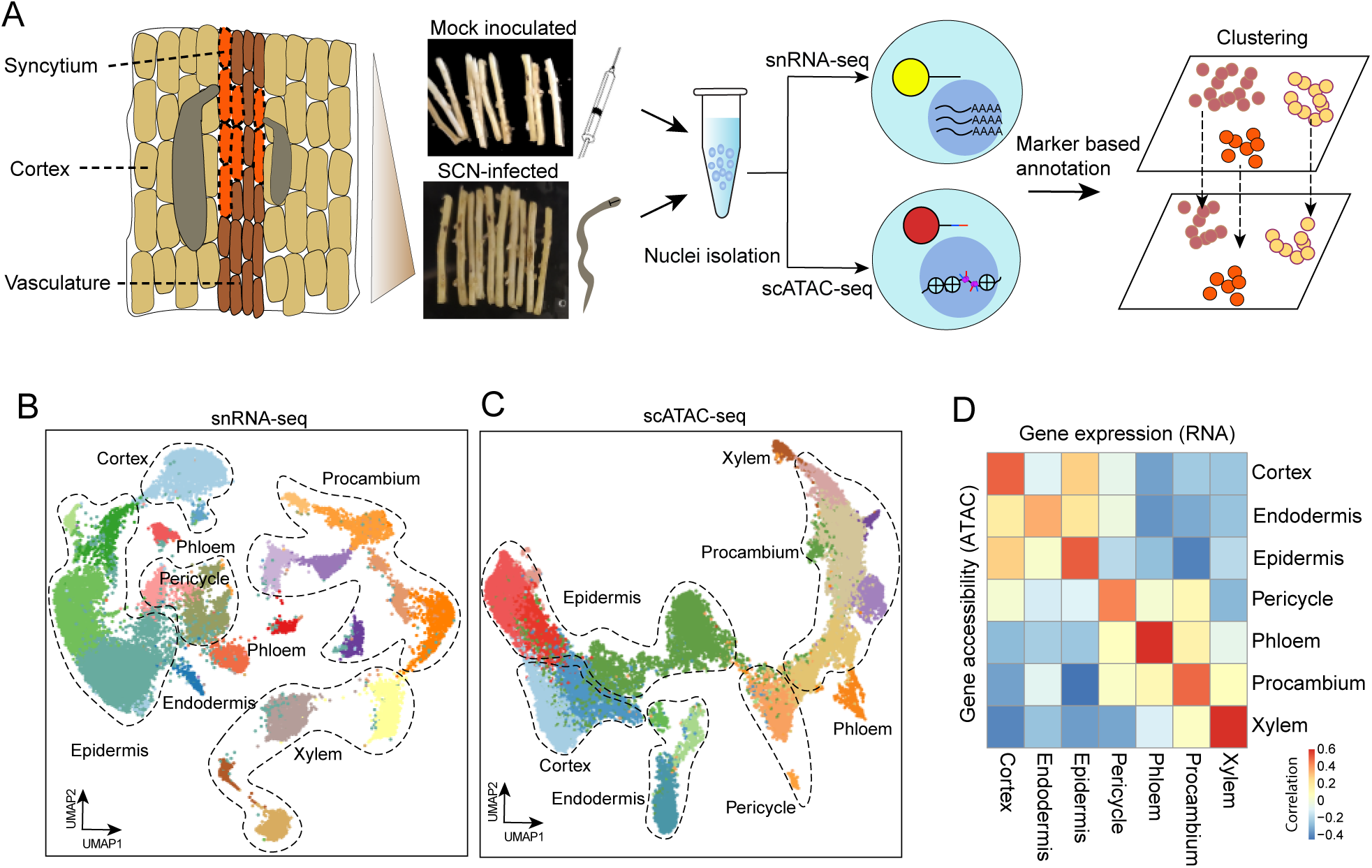
Profiling single-nucleus transcriptomes and chromatin accessibility in SCN-infected soybean roots. (A) Overview of tissue types and experimental design. (B-C) Two-dimensional embeddings using uniform manifold approximation and projection (UMAP) depicting similarity among nuclei based on gene expression (B) and chromatin accessibility (C). (D) Heatmap showing Spearman’s correlation between the 3,000 most variable genes’ expression and accessibility across major cell types.

We performed an integrated analysis of infected and mock-inoculated soybean roots using *Seurat* and *Socrates* workflows [27,28]. This identified 22 snRNA-seq and 18 scATAC-seq clusters, respectively, with consistent nuclei proportions across replicates (Fig. S1D,H; Table S1,S2). To annotate cell clusters, we compiled root marker genes from a published soybean atlas[26]. Cell types were assigned through manual review of marker gene expression and evaluation of enriched biological processes (Fig. S1I,J, Table S3, Methods). Following comprehensive annotation, we identified all major root cell types in both scATAC-seq and snRNA-seq datasets (Fig. 1B,C), with high concordance between gene-body chromatin accessibility (scATAC-seq) and gene expression (snRNA-seq) for corresponding cell types (Fig. 1D). These results demonstrate that our high-quality dataset captures the full diversity of cell types and states in soybean roots.

### Syncytia take on a procambial cell-type signature

During infection, SCN secretes CLAVATA3/EMBRYO SURROUNDING REGION-RELATED (CLE)-like peptide effectors, which mimic endogenous plant CLE peptides. These effectors interact with the CLE receptor TDIF RECEPTOR/PHLOEM INTERCALATED WITH XYLEM (TDR/PXY), activate *WOX4* expression, and promote root cambium proliferation[29]. Consistently, we observed high procambium cell heterogeneity, with notably high expression of two copies of *GmPXY*s and four copies of *GmWOX4*s (Fig. 2A,B). Using classic cell cycle markers, procambium cells were placed into the three canonical cell cycle stages: S stage (Procambium S), M stage (Procambium M), and G2 stage (Procambium G2) (Fig. S2A-C). Additionally, we observed three putative developmental stages: Procambium 1, Procambium 2, and Procambium 3 (Fig. 2A).

**Fig. 2.**
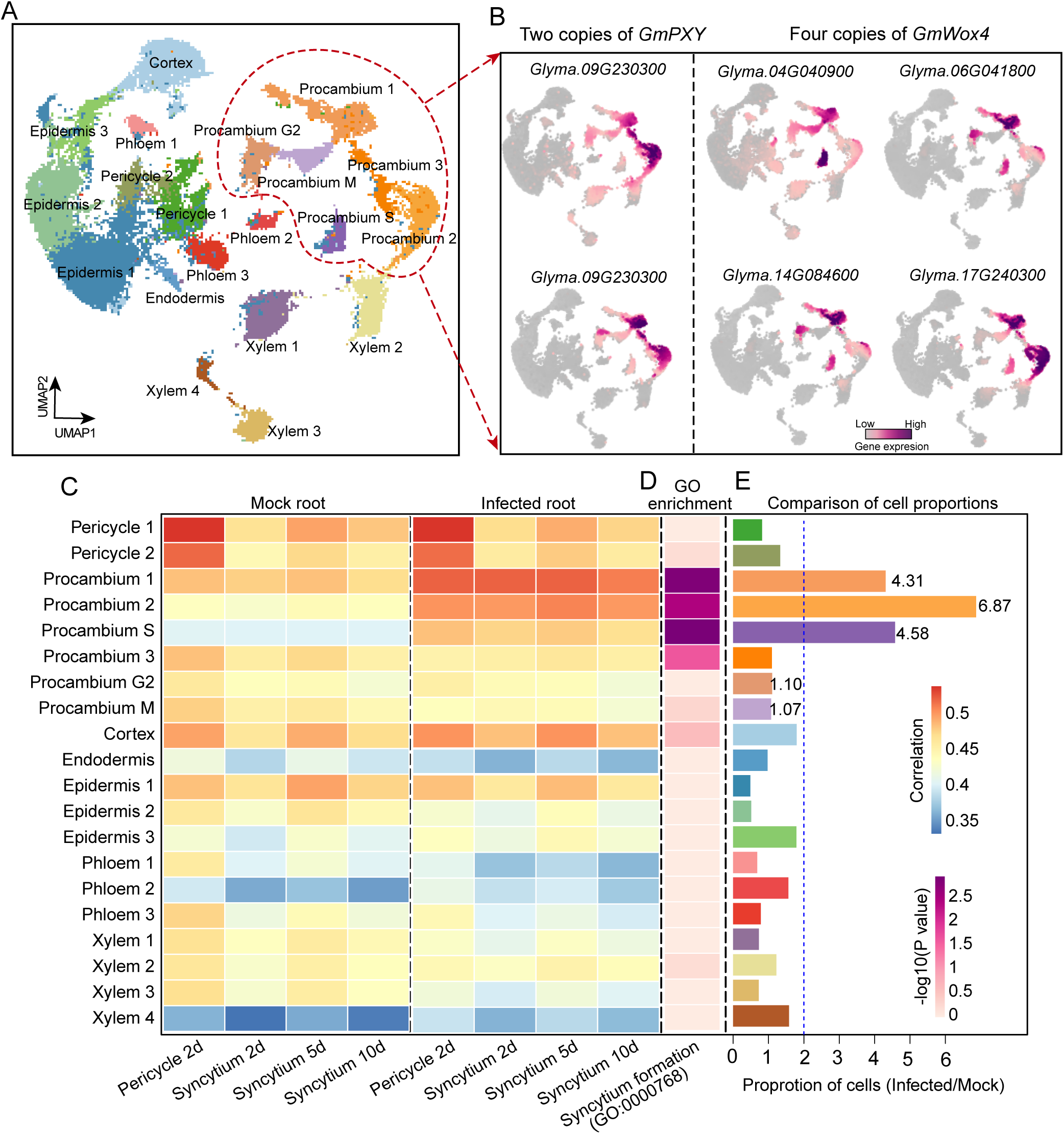
Identification of syncytium-associated procambium cell types. (A) UMAP projection of nuclei from snRNA-seq, colored by cell type. (B) UMAPs showing expression patterns of procambium marker genes *GmPXY*s and *GmWOX4*s. (C) Heatmap showing Spearman’s correlation coefficients between the 3,000 most variable genes from LCM microarray datasets and snRNA-seq cell types for mock (left) and infected (right) roots. (D) GO enrichment analysis across all cell types for the term “Syncytium formation” (GO:0000768). (E) Comparison of cell-type proportions between infected and mock roots.

To determine the syncytium origin, we integrated previous LCM data [15], which included multinucleated syncytia at 2, 5, and 10-DPI, and uninfected root pericycle controls. Correlation analysis showed that the LCM pericycle cells most closely matched the mock-inoculated and SCN-infected pericycle cells, validating the quality of the snRNA-seq data and annotations (Fig. 2C). Interestingly, the three LCM syncytium stages correlated strongly with Procambium 1 and Procambium 2 in SCN-infected roots, whereas this pattern was absent in mock-inoculated roots, which lacked syncytial cells (Fig. 2C). Gene set enrichment analysis (GSEA) further revealed that Procambium 1, Procambium 2, and Procambium S were significantly enriched for the “syncytium formation” category (GO:0000768, P < 0.01; Fig. 2D), whereas other cell types showed no such enrichment. Reflecting infection-specific syncytium cell-state establishment, more than four times as many cells from infected roots compared to mock roots were classified in these three procambium states (Fig. 2E). Endoreduplication, a hallmark of syncytium development, was also evident: the proportion of SCN-infected cells were overrepresented only in the S stage (Procambium S), but not in M or G2 stages, indicating DNA replication without mitotic division, which is a signature of endoreduplication (Fig. 2E). There were no obvious syncytium signatures for Procambium 3, which suggested that it might represent the uninfected procambium cells (Fig. 3C-E). Together, these findings strongly suggest that syncytial cells take on a procambial cell signature. Hereafter, the three cell types, Procambium 1, Procambium 2, and Procambium S, are referred to as syncytium-associated procambium cell types which formed a stronger correlation block than other procambium lineage (Fig. S2D).

**Fig. 3.**
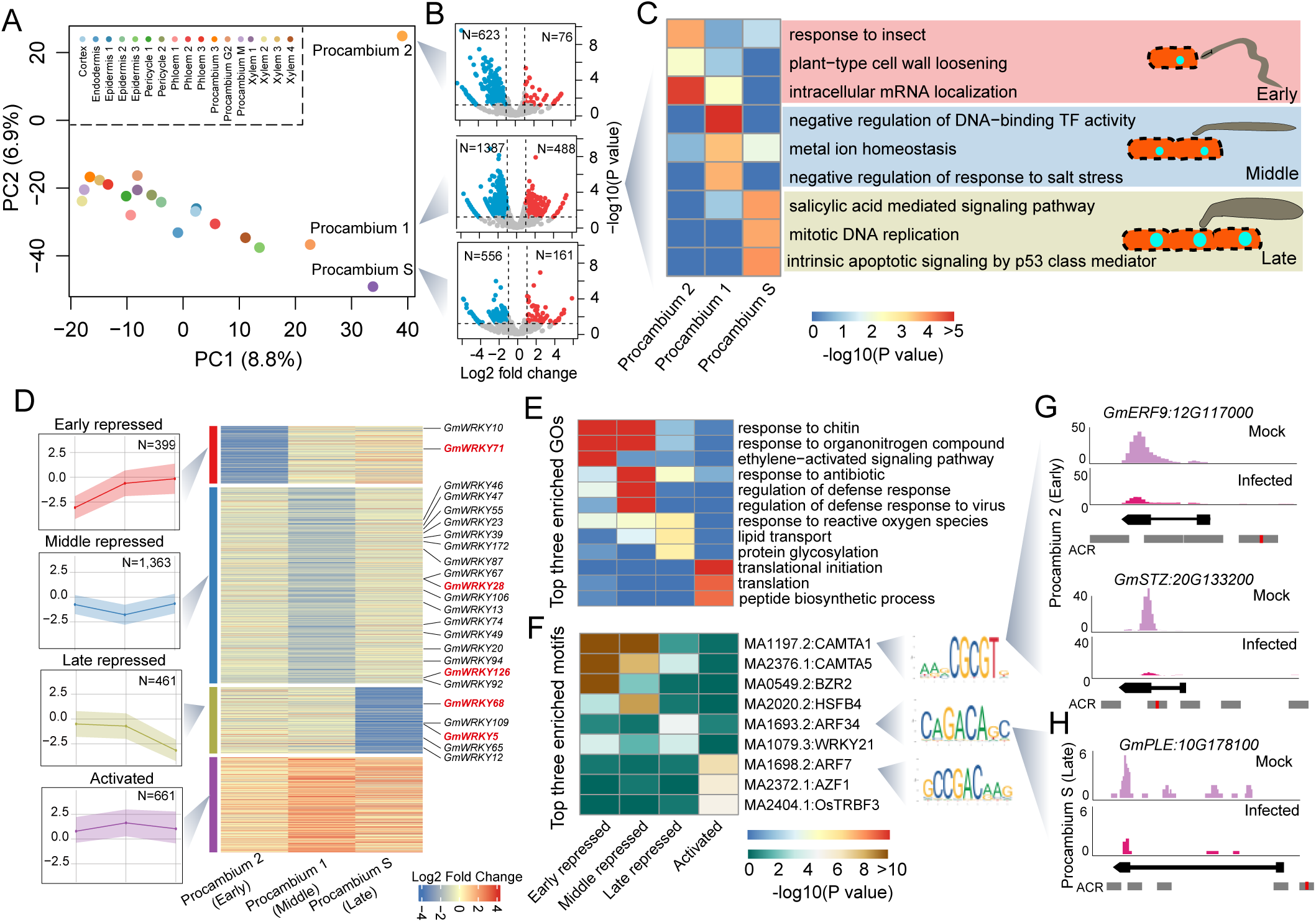
Characterization of differentially expressed genes (DEGs) for syncytium-associated procambium cell types. (A) PCA of log2(Infected/Mock) gene expression changes across all cell types using the 3,000 most variable genes. (B) Volcano plots showing differentially expressed genes (DEGs; fold change > 2, p < 0.05) in the three syncytium-associated procambium cell types: Procambium 2 (top), Procambium 1 (middle), and Procambium S (bottom). (C) Representative top three GO categories enriched in three syncytium-associated procambium cell types. (D) K-means clustering heatmap of DEGs across the three syncytium-associated procambium cell types, identifying four expression groups: Early repressed, Middle repressed, Late repressed, and Activated. *WRKY* genes are labeled at the right; functionally validated genes are highlighted in red. (E) Heatmap showing the three most enriched GO categories for each DEG group. (F) Heatmap of the three most enriched transcription factor motifs within ACRs associated with the four DEG groups. (G) Pseudobulk expression profiles in Procambium 2 (Early) for *GmERF9 (Glyma.12GG117000)* and *GmSTZ (Glyma.20133200)*, both containing CAMTA1 (MA1197.2) motif in their ACRs. (H) Pseudobulk expression coverage in Procambium S (Late) around the *GmPLE (Glyma.10G178100)*.

### Transcriptional reprogramming during syncytium development

To comprehensively characterize transcriptional reprogramming during syncytium development, we first identified differentially expressed genes (DEGs) by comparing cell-types from SCN-infected and mock-inoculated roots (Fig. S3A, Table S4; Methods). Principal component analysis (PCA) based on the fold-change matrix (genes × cell types; Methods) indicated that the three syncytium-associated procambium cell types exhibit a distinct gene regulatory program compared to other cell types (Fig. 3A). Notably, in SCN-infected roots, at least twice as many genes were downregulated as were upregulated in the three syncytium cell types relative to mock-inoculated roots (Fig. 3B). This highlights the relative importance of SCN-mediated repression of host transcription during syncytium formation.

Syncytium formation requires cell fusion at early stages, while endoreduplication becomes more prominent as the syncytium develops into a functional feeding site [30,31]. Exploiting the processes accompanying syncytium development, we ordered the three syncytium-associated procambium cell types according to developmental stage. Accordingly, Gene Ontology (GO) enrichment of upregulated genes reflected the biological processes that accompany syncytium cell development (Fig. 3C): Procambium 2 upregulated genes were enriched for responses to insects and cell wall loosening, reflecting early soybean cell responses to nematode infection; Procambium 1 genes were enriched in negative regulation of DNA-binding transcription factor activity and stress responses, suggesting repression of defense pathways necessary for syncytium development; Procambium S genes were enriched in salicylic acid signaling and mitotic DNA replication, reflecting hormonal responses and endoreduplication [32,33]. Correlation with the LCM dataset, which represents fused multinucleated cells, suggests that Procambium 1 is enriched with multi-nucleated syncytia. Furthermore, we observed a gradient of correlation from uninfected procambium cells (Procambium 3) to Procambium 2, Procambium 1 and Procambium S, indicating a developmental continuum among these states. Accordingly, we refer to Procambium 2, Procambium 1 and Procambium S as early, middle and late developmental stage syncytia, respectively.

To further characterize syncytial transcriptional dynamics, all DEGs from the three syncytium stages were clustered and classified into four gene groups based on expression patterns (Table S5): Early Repressed (*n* = 399), Middle Repressed (*n* = 1,363), Late Repressed (*n* = 461), and Activated (*n* = 661) (Fig. 3D). The three repressed groups were mainly associated with Procambium 2 (early stage), Procambium 1 (middle stage), and Procambium S (late stage) respectively, whereas the Activated group was upregulated across all three stages. GO enrichment of the repressed groups highlighted defense-related pathways, such as responses to chitin, organonitrogen compounds, and reactive oxygen species (ROS), reflecting nematode-mediated suppression of host defenses (Fig. 3E). The WRKY transcription factors play key roles in plant defense, including SCN resistance. Among 174 WRKY genes in soybean [34], 23 were significantly downregulated in the three syncytium cell types in infected roots, suggesting their suppression facilitates syncytia development, whereas their overexpression could potentially enhance resistance (Fig. 3D). Notably, five genes, including *GmWRKY5*, *GmWRKY28*, *GmWRKY68*, *GmWRKY71*, and *GmWRKY126*, were previously validated in hairy root systems [34], to show overexpression inhibited SCN growth and enhanced resistance (Fig. S3D).

To identify key transcription factors regulating the DEGs, we collected accessible chromatin regions (ACRs) associated with each gene group and performed transcription factor motif enrichment analysis (Methods). Motifs for Calmodulin-binding transcription activators (CAMTA1/5, MA1197.2/MA2376.1) [35], heat shock factor HSFB4 (MA2020.2) [36], and WRKY21 (MA1079.3) [34] were enriched in the three repressed gene groups, consistent with their roles in biotic and abiotic stress regulation (Fig. 3F, Table S6). GO analysis of CAMTA1 motif-containing repressed genes revealed enrichment in defense pathways, including response to chitin and ethylene-activated signaling (*P* < 1e-5, Table S7). CAMTA1 targets included transcription factors such as ethylene-responsive element binding factors (*GmERF4* [*Glyma.06G236400, Glyma.12G226600*]), *GmERF9* [*Glyma.12G117000*]) and C2H2-type zinc finger proteins (*GmSTZ* [*Glyma.20G133200*], *GmZF1* [*Glyma.10G257900*], GmRHC2A [*Glyma.13G338400*]) (Fig. 3G, Table S7). Because ERFs and zinc-finger TFs are positive regulators of nematode resistance [37,38], their CAMTA repression likely facilitates syncytium development in susceptible lines.

An auxin response factor (ARF34) motif (MA1693.2) was specifically enriched in the regulatory regions of the Late Repressed group. ARFs are associated with cell cycle regulation and endoreduplication via auxin signaling known to be differentially regulated in developing syncytia [39]. In susceptible soybean, the 5 DPI timepoint encompasses active syncytium expansion, where auxin signaling drives cell cycle modifications and endoreduplication. Consistently, Late Repressed genes with ARF34 motifs were enriched for mitotic cell cycle (GO:0007346) and cytokinesis (GO:0000911) (*P* < 0.05, Table S7). Targets included multiple cell division and endoreduplication-associated genes, such as PLEIADE (GmPLE; Glyma.10G178100), Cyclin-dependent kinase G2 (GmCDKG2; Glyma.11G248500), and Kinesin-like protein for actin-based chloroplast movement 1 (GmKAC1; Glyma.08G279800) (Fig. 3H, Table S7). These findings suggest that endoreduplication in syncytia is likely modulated by ARFs through repression of cell-division–related genes.

In summary, comparative analysis of SCN-infected and mock-inoculated roots defined three molecular stages of early syncytium development. Defense-related genes were strongly repressed, primarily through CAMTA-mediated regulatory networks, facilitating syncytium growth, while repression of cell division-related genes by ARFs supports the endoreduplication process essential for syncytium maturation.

### Chromatin dynamics during syncytium development

To investigate chromatin dynamics underlying gene expression reprogramming, we integrated scATAC-seq and snRNA-seq data for all procambium nuclei from both SCN-infected and mock-inoculated root samples (Fig. 4A, Table S8; Methods). Using snRNA-seq nuclei as a reference, we annotated integrated cell clusters as Early, Middle, Late, and M/G2 stages by tracing their original identities from the snRNA-seq analysis. As before, the Early Stage mainly comprised nuclei from Procambium 2, representing the early response to nematode infection, the Middle Stage nuclei consisted predominantly of Procambium 1, corresponding to multinucleated syncytia formed by cell merging and the Late Stage contained Procambium S nuclei, representing established syncytia undergoing endoreduplication. The M/G2 Stage included nuclei from Procambium M and G2, representing cells in the corresponding cell cycle stages (Fig. 4B). Importantly, gene chromatin accessibility and gene expression showed high correlations across all cell types, reflecting good modality integration (Fig. 4C).

**Fig. 4.**
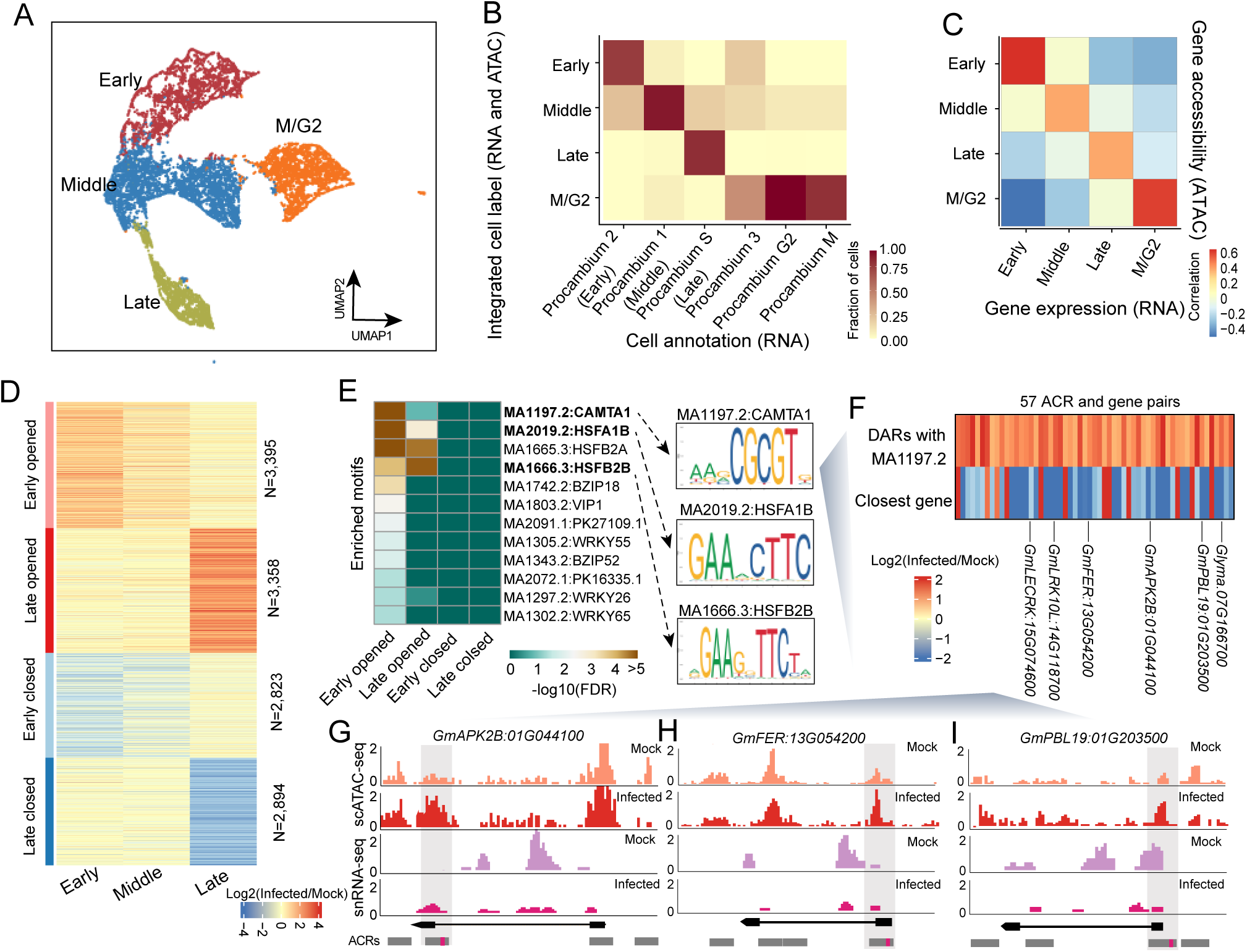
Characterization of differentially accessible regions (DARs) for syncytium-associated procambium cell types. (A) UMAP projection of integrated snRNA-seq and scATAC-seq nuclei from procambium cells, colored by four developmental stages. (B) Overlap between integrated cell-stage labels (rows) and snRNA-seq-derived labels (columns). (C) Heatmap showing Spearman’s correlation between the 3,000 most variable genes’ expression and accessibility across cell stages. (D) K-means clustering heatmap of differentially accessible regions (DARs) across Early, Middle, and Late stages, identifying four DAR groups: Early opened, Late opened, Early closed, and Late closed. Numbers of DARs are indicated at right. (E) Heatmap of all enriched motifs (FDR < 0.05) across the four DAR groups. No enriched transcription factor motifs were detected for the Early closed and Late closed groups. (F) Heatmap of log2(Infected/Mock) changes for DARs containing CAMTA1 (MA1197.2) motifs and their associated DEGs (> two-fold change) in Early and/or Middle stage. Six RLKs related genes were labeled at the bottom. (G-I) Pseudobulk Tn5 integration and gene expression coverage in Early stage (mock vs infected) for RLK genes containing CAMTA1 motif (red dots in ACRs), including *GmAPK2B* (G), *GmFER* (H), *GmPBL19* (I).

Mirroring the snRNA-seq approach, we identified differentially accessible regions (DARs) between SCN-infected and mock-inoculated roots across all syncytium-associated cell types (Table S9, Methods). K-means clustering classified DARs into four developmental groups (Fig. 4D, Table S9): Early Opened (N=3,395), higher chromatin accessibility in Early and/or Middle stages in infected roots; Late Opened (N=3,358), higher chromatin accessibility in Late stage; Early Closed (N=2,823), lower chromatin accessibility in Early and/or Middle stages; Late Closed (N=2,894): lower chromatin accessibility in Late stage. Interestingly, the numbers of opened and closed DARs were comparable across development, in contrast to DEGs, which were predominantly repressed. This highlights the complex relationship between chromatin accessibility and gene expression, influenced by dynamics of transcription factor-transcription factor and transcription factor-DNA interactions.

To identify transcription factor motifs driving these changes, we performed motif enrichment analysis for each DAR group (Methods). CAMTA1 (Calmodulin-binding transcription activator 1, MA1197.2, FDR = 3.53e-14) was most enriched in the Early Opened group, whereas three heat shock factors, including HSFA1B, HSFB2A, HSFB2B, were enriched in both Early and Late Opened groups (Fig. 4E, FDR < 1e-5). CAMTA and HSF family transcription factors play central roles in early biotic and abiotic stress responses [35,40], regulating stress-responsive genes through activation or repression.

Notably, CAMTA1 motifs were enriched in ACRs associated with Early and Middle Repressed DEGs as well as Early Opened DARs, indicating that regulatory regions were accessible whereas gene expression was downregulated. This suggests that CAMTA1 functions as a transcriptional repressor during SCN syncytium development. In Arabidopsis, CAMTA1 and CAMTA3 act redundantly to negatively regulate immunity via multiple hormone signaling pathways [41], though their role in nematode response remains unclear. Analysis of Early Opened DARs containing CAMTA1 motifs and their downstream DEGs revealed that most target genes were downregulated, including six receptor-like kinase (RLK) genes (*GmAPK2B [Glyma.01G044100]*, *GmPBL19 [Glyma.01G203500], Glyma.07G166700, GmFER [Glyma.13G054200], GmLRK10L [Glyma.14G118700], GmLECRK [Glyma.15G074600]*), which are key hubs in plant signaling (Fig. 4F-I). Repression of these RLKs likely suppresses plant cellular processes that facilitate SCN infection and syncytium formation. In summary, integrating single-cell ATAC-seq and RNA-seq data revealed the chromatin dynamics underlying SCN-induced transcriptional reprogramming identifying CAMTA1 as a potential transcriptional repressor that may suppress plant signaling, such as repressing RLKs, during SCN syncytia formation.

### Developmental trajectories reveal syncytial gene regulation dynamics

To investigate chromatin and gene expression dynamics during syncytium development, we performed pseudotime analysis on Early, Middle, and Late stage cells using snRNA-seq nuclei in SCN-infected roots as a reference (Fig. 5A, Methods). Mapping nuclei back to their original snRNA-seq identities confirmed that the trajectory followed the expected developmental order: uninfected cells (Procambium 3) → Early stage (Procambium 2) → Middle stage (Procambium 1) → Late stage (Procambium S) (Fig. 5B). We identified 1,497 genes with significant expression variation along the pseudotime trajectory (Fig. 5C; Methods) and classified them into four temporal expression groups (Stage 1–4). Notably, two copies of *PXY*s (*Glyma.09G230300, Glyma.12G006300*) and one copy of *WOX4* (*Glyma.04G040900*) were upregulated in early development (Fig. S4A, Table S10), consistent with their known roles in early nematode response and promotion of cambium proliferation in the feeding site [29]. These results support the robustness of the syncytium developmental trajectory and its potential for dissecting gene regulatory networks underlying cell identity reprogramming.

**Fig. 5.**
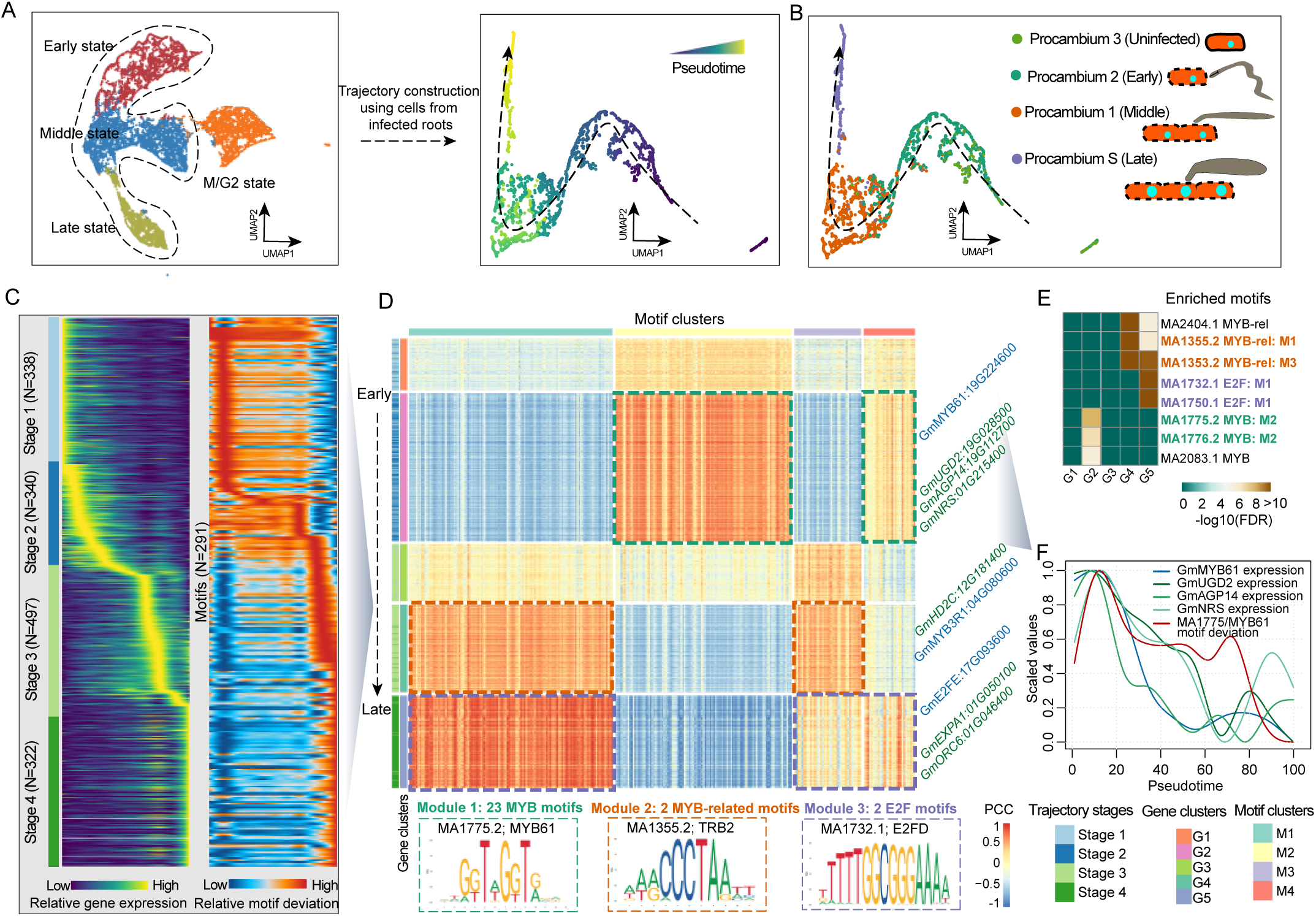
Pseudotime trajectory analysis of syncytium development. (A) Workflow for constructing pseudotime trajectories of syncytium-associated procambium cells. (B) UMAP projection of pseudotime trajectories colored by snRNA-seq-derived cell types. (C) Heatmap showing relative activity of pseudotime-associated genes (left, FDR < 0.05) and motifs (right, FDR < 0.05) along pseudotime. (D) Correlation heatmap between transcription factor motif deviation scores (columns) and pseudotime-associated genes (rows). K-means clustering identified five gene clusters (G1–G5) and four motif clusters (M1–M4). Pearson’s correlation coefficients (PCC) are shown. Three regulatory modules are boxed, with representative motifs shown below. (E) Heatmap showing the three most enriched motifs for ACRs associated with each gene cluster (G1–G5). (F) Expression of *GmMYB61 (Glyma.19G028500)*, MYB61 (MA1775.2) motif deviation, and expression of its putative target genes *GmUGD2 (Glyma.19G028500), GmAGP14 (Glyma.19G112700),* and *GmNRS (Glyma.01G215400)* across the pseudotime trajectory.

To examine transcription factor activity dynamics, we imputed transcription factor motif deviations from scATAC-seq onto snRNA-seq nuclei, identifying 291 transcription factor motifs with pseudotime dynamic chromatin accessibility (Fig. 5C). Correlating transcription factor motif deviations with gene expression revealed five gene clusters (G1–G5 from early to late stages) and four transcription factor/motif clusters (M1–M4) (Fig. 5D). The gene clusters corresponded to the pseudotime stages from early to late stages: Stage 1 → G1, Stage 2 → G2, Stage 3 → G3 & G4, Stage 4 → G5. GO enrichment reflected stage-specific biological processes: early-stage transitions involved metal ion homeostasis and cell wall organization, mid-stage genes supported metabolic reprogramming for syncytium growth, and late-stage genes were associated with chromatin assembly during endoreduplication (Fig. S4B). Integrating gene and transcription factor/motif correlations, we defined three modules: Module 1 (G2:M2&M4), Module 2 (G4:M1&M3), and Module 3 (G5:M1&M4) (Fig. 5D).

Motif enrichment of associated ACRs revealed that MYB, MYB-related, and E2F motifs were significantly enriched in G2 (Stage 2), G4 (Stage 3), and G5 (Stage 4), respectively (Fig. 5E). MYB and MYB-related transcription factors regulate plant defense against multiple pathogens [42,43], whereas E2F transcription factors control cell cycle progression, DNA repair, and endoreduplication, consistent with Stage 4 developmental events. Importantly, enriched motifs were concentrated within high-correlation gene-motif modules: 23 MYB motifs in Module 1, two MYB-related motifs in Module 2, and two E2F motifs in Module 3, whereas none were found outside the modules, indicating that these transcription factors likely act as key activators (Fig. 5D).

Exploring potential regulatory networks, we used module-resolved motif enrichments, cognate motifs near genes and cognate-TF expression to identify putative transcription factors-to-gene target within each module. For example, in Module 1, five MYB family genes (including *GmMYB61*, Glyma.19G224600) were co-expressed with 268 potential targets, including three cell wall modification genes (G*mUGD2, GmAGP14, GmNRS*). In Arabidopsis, *UGD2* is required for cell wall ingrowth in syncytia, and its absence reduces host suitability and syncytium size [44]. Expression patterns of *GmMYB61*, its motif deviation, and target genes were synchronized along pseudotime (Fig. 5F). In Module 2, one MYB-related gene (*GmMYB3R1*) had 218 potential targets, including *GmHD2C*, an HD2-type histone deacetylase potentially repressing defense genes in Stage 3 via histone modification. In Module 3, one E2F family gene (*GmE2FE*) had 136 potential targets, including *GmORC6* (Origin Recognition Complex subunit 6), involved in initiating DNA replication.

In summary, the pseudotime trajectory revealed coordinated dynamics of gene expression and chromatin accessibility during syncytium development and identified key transcription factors, including MYB, MYB-related, and E2F, that likely regulate cell wall modification, chromatin remodeling, and DNA replication.

## Discussion

The spatial and temporal dynamics of plant-microbe interactions have remained challenging to resolve. The study of sedentary endoparasitic nematode-plant interactions is particularly challenging because feeding cells induced by these nematodes are deeply embedded in the vasculature of the root and are therefore only represented by a small fraction of cells relative to the total root cell population. However, single-cell technologies offer unprecedented resolution to study these interactions. The primary goal of this study was to apply single-cell multi-omics technology to SCN-infected soybean roots to gain new insights on the molecular basis of syncytia initiation and development. We performed snRNA-seq and scATAC-seq on 56,448 high-quality nuclei from infected and mock-inoculated roots, generating the first multi-omic atlas of soybean SCN root responses. Comparative transcriptomics and epigenomics identified multiple syncytial developmental stages with reprogrammed procambial molecular signatures, consistent with a recent study[45]. Beyond clarifying the cellular signature of SCN syncytia, this developmental trajectory provided insights into the ordinal molecular events behind syncytia formation. To facilitate further discovery, we have incorporated this dataset into our soybean multi-omic atlas database (https://soybean-atlas.com/) [26], which includes chromatin accessibility and gene expression across multiple tissues and developmental stages. The interactive platform allows researchers to explore gene regulatory network changes response to SCN at cell-type resolution.

Identifying syncytium-specific gene programs provides insights into both plant defense and nematode virulence, that can inform targeted breeding efforts to reduce yield losses. Here, we identified 34 potential marker genes significantly enriched in syncytia cells relative to other cell types in SCN-infected and mock-inoculated roots. These markers acted across multiple developmental stages: early response (2 genes), multinucleated syncytia formed by cell fusion (25 genes), and endoreduplicated syncytia (7 genes) (Table S11). Integrating scATAC-seq identified candidate syncytium-specific *cis*-regulatory regions, providing a valuable resource to understand syncytium development at the transcriptome level and to clone syncytium-specific regulatory regions for developing SCN resistant soybean using genetic engineering or synthetic biology approaches. For example, these regulatory regions can drive targeted expression of transgenes in syncytia, enabling precise defense activation where the nematode feeds, minimizing off-target effects on plant growth [46].

Additionally, we revealed CAMTA and MYB transcription factors as putative syncytium establishment regulators. CAMTA represses multiple immunity-related gene expression, enabling syncytium development (Fig. 4E-F). Later in development, MYB upregulates gene expression underpinning cell wall modification, neighboring cell fusion and syncytium expansion (Fig. 5D). Future genome editing efforts targeting TFs or their target motifs will clarify their roles in syncytium formation and potentially improve SCN resistance in soybean. These datasets significantly advance our understanding of syncytium reprogramming and provide a resource for reconstructing gene regulatory networks underlying its development.

Although this study characterized SCN syncytium development in a susceptible cultivar, we observed strong downregulation of defense-related genes, highlighting their role in soybean SCN response. Specifically, we identified 23 *WRKY* genes downregulated during syncytium development. WRKY genes are known mediators of SCN resistance, with five of the DEGs (*GmWRKY5, GmWRKY28, GmWRKY68, GmWRKY71,* and *GmWRKY128*) previously enhancing SCN resistance when overexpressed in an SCN-susceptible soybean background [34] (Fig. S3D). Notably, *GmWRKY28* is located within the *qSCN3-1* QTL [47], providing genetic evidence for its resistance function (Fig. S3B-D). In this way, we show how our data can integrate with QTL, GWAS, and orthologous resistance genes to identify candidates (Table S5) for improving SCN resistance via genetic engineering (Table S5).[46]

This work establishes a cell-type-resolved framework for characterizing soybean-nematode regulatory interactions, which can be extended to other plant-nematode systems. Future studies applying this single-cell framework to soybean genotypes with varying SCN resistance, at multiple infection time points, will further advance our understanding of the molecular mechanisms underlying host-pathogen dynamics and facilitate the development of nematode-resistant cultivars.

## Methods

### Plant materials

The SCN (*Heterodera glycines* Ichinohe) inbred population PA3 (HG type 0) population used in this study was maintained on Williams 82 in the greenhouse at the University of Georgia. The soybean cultivar Williams 82 to was used for infection assays.

### Nematode infection assays and tissue collection

Nematode infection assays were conducted as described earlier [15]. Briefly, soybean seeds were germinated in ragdolls for three days at 28°C in the dark. Seedlings with uniform radicles were selected for SCN inoculation. Approximately 300 second-stage juveniles (J2) suspended in 0.08% agarose were applied 1 cm above the root tip of each soybean root. Mock-inoculated seedlings were treated with 0.08% agarose without nematodes. The infection was synchronized by washing roots 24 hours after inoculation with running tap water to remove J2s outside the root. Mock-inoculated seedlings were treated the same way. Washed seedlings were rolled into ragdolls, placed in Hoagland’s nutrient solution with constant aeration, and grown in a plant growth chamber at 26°C with a photoperiod of 16 h of light and 8 h of dark for four more days. Five days post-inoculation, root pieces of approximately 1-1.5 cm flanking the inoculation point were excised from both SCN-infected and mock-inoculated samples. Two biological replicates, each consisting of 20 root pieces, were used for nuclei extraction.

### scATAC-seq library preparation

Nuclei were isolated and purified following the protocol described previously [26], with minor modifications. Briefly, root tissues were finely chopped with single edge razor blades on ice for approximately 2 minutes in 600 μL of pre-chilled Nuclei Isolation Buffer (NIB; 10 mM MES-KOH, pH 5.4; 10 mM NaCl; 250 mM sucrose; 0.1 mM spermine; 0.5 mM spermidine; 1 mM DTT; 1% BSA; and 0.5% Triton X-100). The homogenate was passed through a 40-μm cell strainer and centrifuged at 500 × g for 5 minutes at 4 °C. The supernatant was discarded, and the pellet was gently resuspended in 500 μL of NIB wash buffer (10 mM MES-KOH, pH 5.4; 10 mM NaCl; 250 mM sucrose; 0.1 mM spermine; 0.5 mM spermidine; 1 mM DTT; and 1% BSA). The suspension was filtered through a 40-μm cell strainer and carefully layered onto 1 mL of 35% Percoll (prepared by mixing 35% Percoll with 65% NIB wash buffer) in a 1.5-mL centrifuge tube. Nuclei were enriched by centrifugation at 500 × g for 10 minutes at 4 °C. Following centrifugation, the supernatant was removed, and the nuclei pellet was resuspended in 10 μL of diluted nuclei buffer (DNB; 10x Genomics, Cat# 2000207). For quality control, 5 μL of the nuclei suspension were diluted tenfold, stained with DAPI (Sigma, Cat# D9542), and examined under a fluorescence microscope using a hemocytometer to assess integrity and concentration. The nuclei concentration was then adjusted to 3,200 nuclei/μL using DNB buffer, and 5 μL (approximately 16,000 nuclei) were used as input for single-cell ATAC-seq library construction.

Single-cell ATAC-seq libraries were generated using the Chromium Next GEM Single Cell ATAC v1.1 kit (10x Genomics, Cat# 1000175) according to the manufacturer’s protocol (Chromium Next GEM Single Cell ATAC Reagent Kits v1.1 User Guide, CG000209 Rev E). The resulting libraries were sequenced on an Illumina NovaSeq 6000 platform using dual-index mode with 8 cycles for the i7 index and 16 cycles for the i5 index.

### snRNA-seq library preparation

Nuclei isolation and purification for snRNA-seq were adapted from the previously described scATAC-seq procedure with modifications to minimize RNA degradation and leakage. Briefly, root tissues were finely chopped on ice for approximately 1 minute in 600 μL of pre-chilled Nuclei Isolation Buffer (NIB; 10 mM MES-KOH, pH 5.4; 10 mM NaCl; 250 mM sucrose; 0.4 U/μL RNase inhibitor [Roche, Protector RNase Inhibitor, Cat# RNAINH-RO]; 0.1% NP-40; and 0.5% BSA). The homogenate was passed through a 40-μm cell strainer and centrifuged at 500 × *g* for 5 minutes at 4 °C. The supernatant was removed, and the pellet was resuspended in 500 μL of NIB wash buffer (10 mM MES-KOH, pH 5.4; 10 mM NaCl; 250 mM sucrose; 0.5% BSA; and 0.2 U/μL RNase inhibitor). The suspension was then filtered through a 10-μm cell strainer and carefully layered onto 1 mL of 35% Percoll (prepared by mixing 35% Percoll with 65% NIB wash buffer) in a 1.5-mL centrifuge tube. Nuclei were enriched by centrifugation at 500 × *g* for 10 minutes at 4 °C.

After centrifugation, the supernatant was carefully removed, and the pellet was resuspended in 50 μL of NIB wash buffer. To assess nuclei quality and concentration, 5 μL of the nuclei suspension were diluted tenfold, stained with DAPI (Sigma, Cat# D9542), and examined under a fluorescence microscope using a hemocytometer. The nuclei concentration was then adjusted to 1,000 nuclei/μL with diluted nuclei buffer (DNB; 10x Genomics, Cat# 2000207). A total of approximately 16,000 nuclei were used as input for snRNA-seq library construction.

snRNA-seq libraries were generated using the Chromium Next GEM Single Cell 3′ GEM Kit v3.1 (10x Genomics, Cat# PN-1000123) following the manufacturer’s protocol (Chromium Next GEM Single Cell 3′ Gene Expression v3.1 Dual Index User Guide, CG000315 Rev B). The resulting libraries were sequenced on an Illumina NovaSeq 6000 platform in dual-index mode with 10 cycles each for the i7 and i5 indices.

### scATAC-seq raw data processing and clustering

Processing of raw scATAC-seq data followed the general workflow described previously [26], with minor adjustments for the current study. Briefly, raw BCL files were demultiplexed and converted into FASTQ format using the 10X Genomics tool Cell Ranger ATAC (cellranger-atac makefastq, v1.2.0) with default parameters. Primary read processing, including adapter and quality trimming, alignment, and barcode attachment/correction, was performed using cellranger-atac count (v1.2.0). Reads were mapped to a composite reference consisting of the soybean Williams 82 v4 genome, the *Glycine max* organelle genomes (NCBI Reference Sequences: NC_007942.1 and NC_020455.1), and the soybean cyst nematode (*Heterodera glycines*) genome (SCNBase, TN10 strain). Properly paired, uniquely mapped reads with mapping quality scores greater than 30 were retained using samtools view (v1.6; parameters: -f 3 -q 30) [48]. Reads containing alternative alignments (XA tags) were also retained to minimize bias from the ancient whole-genome duplication events in soybean. PCR duplicates were removed on a per-nucleus basis using Picard (http://broadinstitute.github.io/picard) MarkDuplicates (v2.16; BARCODE_TAG=CB REMOVE_DUPLICATES=TRUE). Reads aligning to mitochondrial, chloroplast were quantified per barcode and excluded from downstream analyses. Due to their low abundance, reads mapping to the nematode genome were also excluded from all downstream analyses.

To eliminate artifacts, alignments overlapping regions with excessive Tn5 integration bias or potential collapsed sequences were removed. Blacklisted regions were defined as 1-kb windows exhibiting greater than fourfold coverage over the genome-wide median in either Tn5-treated genomic DNA or ChIP-seq input controls. Following filtering, BAM alignments were converted into single-base Tn5 insertion sites in BED format by shifting read coordinates by +4 bp and −5 bp for positive and negative strands, respectively, and retaining only unique integration sites per nucleus.

Nuclei identification and quality control were performed using the R package Socrates [27]. The BED file containing single-base Tn5 integration sites was imported along with the soybean genome annotation (Phytozome, Gmax_508_Wm82.a4.v1). Accessible chromatin regions (ACRs) were called using the callACRs function with the following parameters: genome.size = 8.0e8, shift = −75, extsize = 150, and FDR = 0.1. This step provided an estimate of the fraction of Tn5 integration sites located within ACRs for each nucleus. Per-nucleus metadata were generated using the buildMetaData function with a transcription start site (TSS) window size of 1 kb (tss.window = 1000). Sparse binary matrices were constructed using generateMatrix with a 500-bp window size. High-quality nuclei were retained based on the following thresholds: ≥1,000 unique Tn5 insertion sites, ≥20% of insertions within 2 kb of TSSs, ≥20% of insertions within ACRs, and ≤20% of insertions mapped to organelle genomes.

Integrated clustering across samples and biological replicates was conducted using Socrates [27]. Low-quality features and nuclei were filtered out using the cleanData function (min.c = 100, min.t = 0.01), removing windows accessible in <1% of nuclei and nuclei with <100 accessible ACRs. The resulting nucleus-by-window matrix was normalized using the term frequency–inverse document frequency (TF–IDF) algorithm with L2 normalization (doL2 = TRUE). Dimensionality reduction was performed using reduceDims (method = “SVD”, n.pcs = 25, cor.max = 0.4) to remove components correlated with sequencing depth. The reduced embedding was visualized with UMAP (projectUMAP, k.near = 15). Putative doublets (∼5% of nuclei) were identified and removed by applying a modified Socrates workflow on individual libraries using the detectDoublets and filterDoublets functions (filterRatio = 1.0, removeDoublets = TRUE). To correct for batch effects, we applied the Harmony algorithm with non-default settings (do_pca = FALSE, vars_use = c(“batch”), tau = 5, lambda = 0.1, nclust = 50, max.iter.cluster = 100, max.iter.harmony = 50). The nuclei embeddings were further reduced using UMAP (metric = “correlation”, k.near = 15), and final clustering was performed using callClusters (res = 0.8, k.near = 15, cl.method = 3, m.clust = 25, min.reads = 5e5).

### snRNA-seq raw data processing and clustering

Raw snRNA-seq reads were processed using STARSolo [49] for alignment and gene-level quantification against a combined reference genome consisting of the soybean Williams 82 v4 assembly and the soybean cyst nematode (*Heterodera glycines*, TN10 strain; SCNBase). The following parameters were applied in STARSolo to filter UMIs, remove empty droplets, and count multi-mapping reads: --soloUMIfiltering MultiGeneUMI_CR and --soloCellFilter EmptyDrops_CR. Only uniquely mapped reads were retained for downstream analyses.

Filtered expression matrices were processed using the Seurat R package (v4) [28]. Low-quality nuclei and empty droplets were excluded based on multiple quality metrics. Specifically, nuclei were removed if their detected gene counts fell below the median gene count minus two times the median absolute deviation, or if their total UMI counts were below the lower 10th percentile. Nuclei with more than 15% of total reads mapping to organelle genes were also excluded. The filtered datasets were normalized using SCTransform, followed by principal component analysis (PCA) via the RunPCA function. Putative doublets were detected and removed using the DoubletFinder R package [50].

To mitigate the influence of cell cycle heterogeneity on clustering, cell cycle scores were calculated for each nucleus using Seurat’s CellCycleScoring function. Soybean cell cycle–associated genes were identified based on their orthologs in *Arabidopsis thaliana* as reported previously [51]. The effects of cell cycle variation were regressed out during normalization using SCTransform with the parameter vars.to.regress.

Two biological replicates were prepared for each library and integrated using the Harmony R package [52] to correct for batch effects. The integrated dataset was further analyzed using Seurat’s dimensional reduction and clustering functions: UMAP embedding was generated with RunUMAP(reduction = “harmony”, dims = 1:25), the nearest-neighbor graph was computed with FindNeighbors(reduction = “harmony”, dims = 1:25), and clusters were identified using FindClusters(resolution = 0.5).

### Cell-type annotation for snRNA-seq

Cell-type identities were assigned to each cluster using a combination of marker gene-based annotation and gene set enrichment analysis [53]. First, a curated list of cell-type-specific marker genes known to localize to discrete root cell types or tissue domains was compiled based on our previous study [26] (Table S3). For each cluster, gene expression was quantified by summing UMI counts within gene bodies and aggregating all nuclei belonging to the cluster. The resulting raw count matrix was normalized using the cpm function in *edgeR* [54], and *Z*-scores were computed for each marker gene across all clusters using the scale function in R. Cell identities were assigned based on clusters showing the highest *Z*-scores for well-characterized marker genes. Ambiguous clusters exhibiting similar expression patterns to established cell types were grouped with those key cell types, reflecting potential state-level variation within the same lineage (Fig. S3). To improve visualization of gene expression patterns, normalized expression values were smoothed by constructing a diffusion nearest-neighbor graph.

To identify the syncytium cell type, the CPM-normalized expression matrix was compared against laser capture microdissection (LCM) microarray data distinguishing pericycle and syncytium tissues [15]. Only probes uniquely mapped to a single gene (*n* = 13,025) were used for this correlation-based comparison. This approach confirmed our cluster annotations and identified a subset of procambium-derived cells as the syncytium population.

Gene set enrichment analysis (GSEA) [53] was performed using the R package fgsea, following a previously described approach. Briefly, a reference panel was generated by uniformly sampling nuclei from each cluster, with the total number of sampled nuclei set to the average cluster size. For each cluster, UMI counts were aggregated across nuclei, and differential expression analysis was conducted between each cluster and the reference panel using edgeR [54]. Ranked gene lists, sorted by log2 fold change, were then used as input for fgsea. Gene ontology (GO) terms containing fewer than 10 or more than 600 genes were excluded. GO categories were considered significantly enriched at a false discovery rate (FDR) < 0.05 based on 10,000 permutations. Cell-type assignments were further supported by examining the top enriched GO terms consistent with the expected biological functions of each cell type.

### Cell-type annotation for scATAC-seq

Cell-type annotation for scATAC-seq data followed the same general strategy used for snRNA-seq, with minor adjustments specific to chromatin accessibility profiling. Instead of gene expression, we calculated gene-level chromatin accessibility scores based on the number of Tn5 integration sites within each gene body, including 500 bp upstream and 100 bp downstream regions. The resulting raw counts were normalized using the *cpm* function in edgeR [54]. Cell types were assigned to each cluster following the same procedure used for snRNA-seq, including evaluation of marker gene chromatin accessibility patterns and GO term enrichment profiles to confirm biological consistency.

To assess cross-modality concordance, we calculated the Pearson correlation coefficients between chromatin accessibility scores and gene expression levels for the top 1,000 variable genes identified in the snRNA-seq dataset. Correlation values ranged from 0.4 to 0.7 across corresponding cell types in the two modalities, consistent with results reported in previous studies [26,27] (Fig. 1D).

### Accessible chromatin region (ACR) identification

Following cell clustering and annotation, accessible chromatin regions (ACRs) were identified for each cluster using all Tn5 integration sites. Peak calling was performed with MACS2 [55] using non-default parameters (--extsize 150 --shift -75 --nomodel --keep-dup all). To mitigate potential bias associated with differences in sequencing depth across clusters, we adjusted the *q*-value thresholds based on the total number of Tn5 integration events per cell type as follows: for fewer than 10 million integrations, --qvalue 0.1; for 10–25 million, 0.05; for 25–50 million, 0.025; for 50–100 million, 0.01; and for more than 100 million, 0.001.

Each peak was refined to a 500-bp window centered on its coverage summit. To generate a comprehensive reference set, peaks from all clusters were merged into a unified master peak list using a custom script. Chromatin accessibility scores for each peak were quantified based on Tn5 integration counts and normalized using the *cpm* function in edgeR [54]. Peaks with normalized accessibility values below 4 CPM across all cell types were excluded from downstream analyses.

### Identification of differentially expressed genes (DEGs)

To identify DEGs between SCN-infected and mock-inoculated roots within each cell type, a reference panel was first constructed by uniformly sampling nuclei from both conditions, with the total number of nuclei set to 90% of the smaller sample size. UMI counts were then aggregated across nuclei within each cell type for each replicate. Differential expression analysis was performed for all expressed genes (CPM > 0.1) using the *edgeR* R package [54]. To reduce the impact of random sampling, this procedure was repeated 1,000 times. Genes exhibiting a median fold change > 2 and a *p*-value < 0.05 across iterations were considered DEGs.

### Identification of differentially accessible regions (DARs)

DARs were identified using an analogous approach. A reference panel was generated by uniformly sampling nuclei from SCN-infected and mock-inoculated roots, again using 90% of the smaller sample size. Tn5 integration counts were aggregated across nuclei within each cell type for each replicate. Differential accessibility analysis was conducted for all peaks with CPM > 4 using edgeR [54]. To account for random sampling effects, this analysis was repeated 1,000 times. Peaks with a median fold change > 2 and *p*-value < 0.05 were designated as DARs.

### *De novo* syncytium marker identification

*De novo* marker genes for syncytium cell clusters were identified using the FindAllMarkers function in Seurat [28] with the following parameters: test.use = “wilcox”, logfc.threshold = 1, only.pos = TRUE, and min.pct = 0.1. The resulting markers were further filtered to retain only those with an adjusted *p*-value < 1 × 10⁻⁵ and log2 fold change > 2. To define the final syncytium-specific markers, these filtered markers were intersected with genes significantly upregulated in the infected root compared to mock controls (DEGs) using a stringent threshold of log₂ fold change > 2.

### Transcription factor motif deviation score calculation

Transcription factor motif deviation scores for individual nuclei were computed using chromVAR [56], with the non-redundant core plant position weight matrix (PWM) database from JASPAR2022 [57]. The input matrix was filtered to retain only ACRs containing at least 10 fragments and nuclei with a minimum of 100 accessible ACRs. Bias-corrected motif deviation scores were smoothed across nuclei and integrated into the UMAP embedding for visualization, following the same approach used for visualizing gene body chromatin accessibility.

### Transcription factor motif enrichment

Transcription factor motif occurrences in all ACRs were first identified using FIMO [58] from the MEME Suite with position weight matrices (PWMs) from the non-redundant core plant motif database in JASPAR2024 [57]. To evaluate motif enrichment in cell-type-specific ACRs (ctACRs), the motif distribution in each ctACR set was compared against a control set of “constitutive” ACRs, defined as regions with minimal variability across cell types (fold change < 2 and p > 0.1). Enrichment for each motif in each cell type was assessed using Fisher’s exact test (alternative = “greater”). Multiple testing was controlled using the Benjamini-Hochberg method, and motifs with FDR < 0.05 were considered significantly enriched in cell-type-specific ACRs. Similarly, motif enrichment in activating versus repressing ACRs was tested using Fisher’s exact test, with motifs showing p < 0.01 considered significantly enriched.

### Integration of procambium scATAC-seq and snRNA-seq datasets

Procambium nuclei from scATAC-seq and snRNA-seq datasets were integrated using the uiNMF workflow implemented in the liger R package [59]. Three matrices, including nuclei x gene accessibility, nuclei x ACR and nuclei x gene expression, were first extracted to retain only procambium nuclei based on prior clustering results. Integration was performed using the unshared feature iNMF workflow, which leverages gene body chromatin accessibility, exhibiting positive correlation with gene expression during non-negative matrix factorization. Unshared features (ACRs) were used solely to capture relationships among scATAC-seq nuclei, preserving local neighborhoods in the co-embedding, while repressive ACRs contributed minimally.

The nuclei x ACR matrix was normalized using TF-IDF (via Socrates [27]) followed by the normalize function of liger with default parameters. The matrix was then rescaled such that the sum of all accessible regions per nucleus equaled 1. Using the Seurat framework, the top 2,000 most variable features were identified with FindVariableFeatures (selection.method = “vst”, nfeatures = 2000). The normalized nuclei × ACR matrix was scaled using scaleNotCenter and stored as the set of unshared features for downstream integration.

For shared features (gene IDs) between scATAC-seq and snRNA-seq, genes within the inner 98% quantile of each modality were selected, and the intersecting set was retained. Nuclei × gene activity and nuclei × gene expression matrices were normalized using default settings. Variable genes were selected using selectGenes with var.thresh = 0.1, datasets.use = “RNA”, unshared = TRUE, unshared.datasets = list(2), and unshared.thresh = 0.2. Normalized matrices were scaled using scaleNotCenter. Integration was then performed with optimizeALS (k = 30, use.unshared = TRUE, max_iters = 30, thresh = 1e-10), followed by quantile normalization (quantile_norm) using snRNA-seq as the reference dataset.

To impute scATAC-seq modalities onto the snRNA-seq nuclei, the imputeKNN function from liger [59] was applied. This approach estimates motif deviation scores and normalized chromatin accessibility values from scATAC-seq nuclei for individual snRNA-seq barcodes, producing integrated data containing gene expression, chromatin accessibility, and motif deviation information for each nucleus.

### Pseudotime trajectory analysis

Pseudotime trajectories were constructed following a previously published approach. Specifically, the calcPseudo function from the associated GitHub repository (https://github.com/plantformatics/maize_single_cell_cis_regulatory_atlas) was used with parameters cell.dist1 = 0.95 and cell.dist2 = 0.95, generating pseudotime estimates for individual nuclei along specific developmental branches. Genes exhibiting significant expression variation along each trajectory were identified using the sigPseudo2 function from the same repository (FDR < 0.05). For visualization, expression profiles of significant genes were smoothed using generalized additive models applied to 500 equally spaced pseudotime bins, as described previously [27].

To link transcription factors (TFs) with gene expression dynamics across pseudotime, Pearson correlation analysis was performed between TF motif deviation scores and genes with significant pseudotime variance. Gene and TF/motif clusters were then determined using k-means clustering, with *k* = 5 for genes and *k* = 4 for motifs. The optimal number of clusters was chosen based on the elbow and silhouette methods.

## Data availability

All datasets generated in this study are available at GEO (Accession number: GSE310956). All original code has been deposited at Github (https://github.com/schmitzlab/Gm_SCN_infected_root).

We add the data in this study to our previous web soybean multi-omic atlas (https://soybean-atlas.com/). Any additional information required to reanalyze the data reported in this paper is available from the lead contact upon request.

## Acknowledgement

This research was supported by the United Soybean Board (2432-201-0102 to RJS and MGM), the National Science Foundation (IOS-1856627 to RJS) and the University of Georgia Office of Research (to RJS). This study was supported in part by resources and technical expertise from the Georgia Advanced Computing Resource Center, a partnership between the University of Georgia’s Office of Research and Office of the Vice President for Information Technology. We would like to thank Yinxin Dong and Yangyang Xu for their assistance in maintaining an organized laboratory environment and Dean Kemp and Kurk Lance for SCN population maintenance.

## Author contributions

R.J.S., X.Z. and M.G.M designed the research. X.Z., Z.L., V.A.G. performed the experiments. X.Z., X.L., H.Z., M.A.M., M.G.M. and R.J.S. analyzed the data. X.Z., V.A.G., and R.J.S. wrote the manuscript. The authors read and approved the final manuscript.

