## Supplementary material for "Single-cell multi-omic characterization of the soybean root response to cyst nematode infection": Figure S1

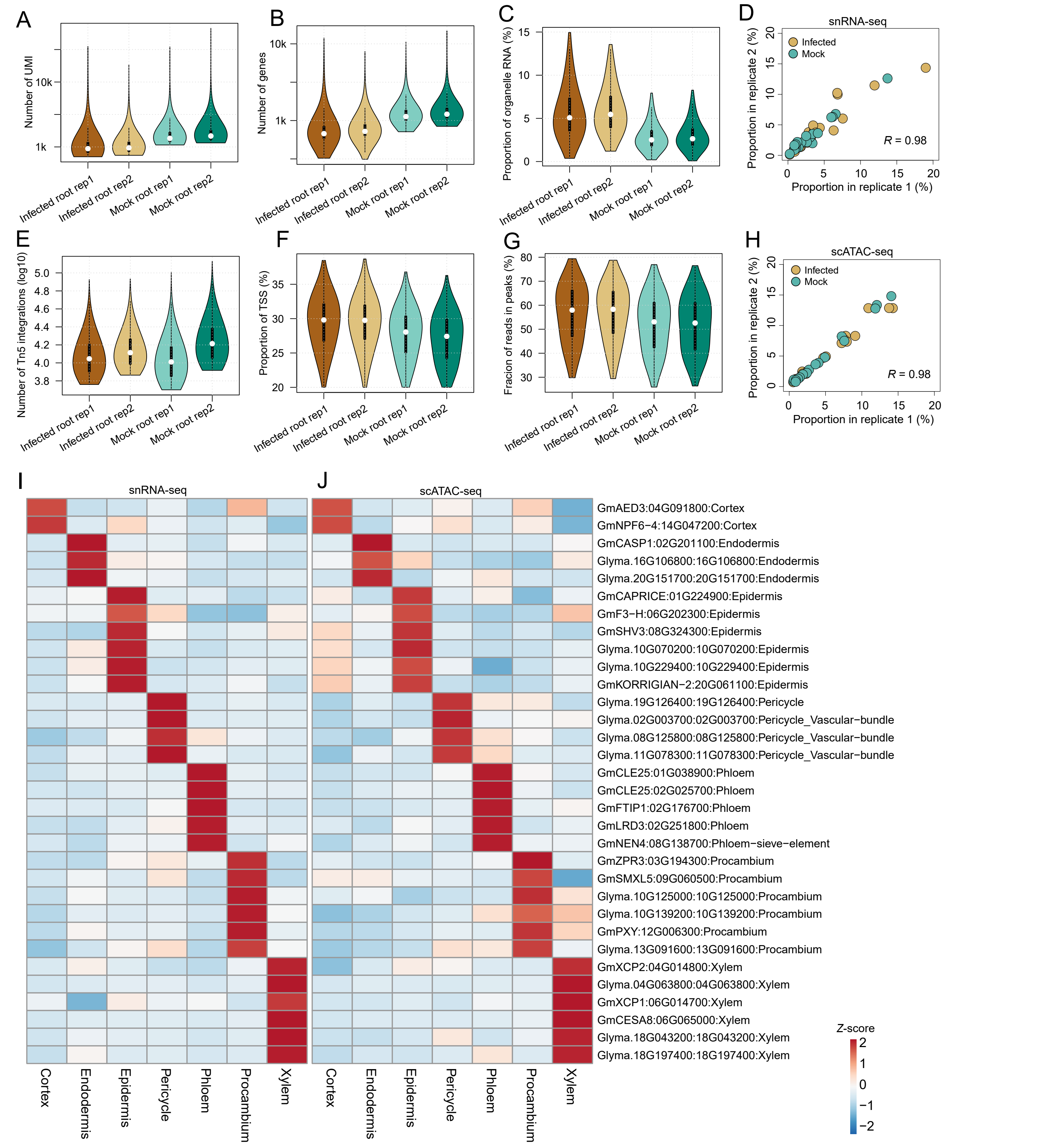


**Fig. S1 Quality control of soybean root snRNA-seq and scATAC-seq datasets**

(A-D) Quality control of snRNA-seq: distribution of total number of UMIs (A); distribution of number of detected genes (B); distribution of the proportion of reads from organelles (C); and correlation of cell proportions between the two replicates across snRNA-seq clusters for mock and infected root (Pearson's correlation coefficient: 0.98) (D).

(E-H) Quality control of scATAC-seq: distribution of Tn5 integration sites per nucleus across the four samples. (E); distributions of the proportion of Tn5 integration sites within the promoter regions, encompassing the 1-kb flanking regions around gene transcription start sites (TSSs) (F); distributions of the proportion of Tn5 integration sites within peaks per nucleus (G); and Correlation of cell proportions between the two replicates across scATAC-seq clusters for mock and infected root (Pearson's correlation coefficient: 0.98) (H).

(I-J) Z score heatmaps showing gene expression (I) and gene chromatin accessibility (J) and for representative marker genes across major cell types in soybean roots.
