## Supplementary material for "Single-cell multi-omic characterization of the soybean root response to cyst nematode infection": Figure S2

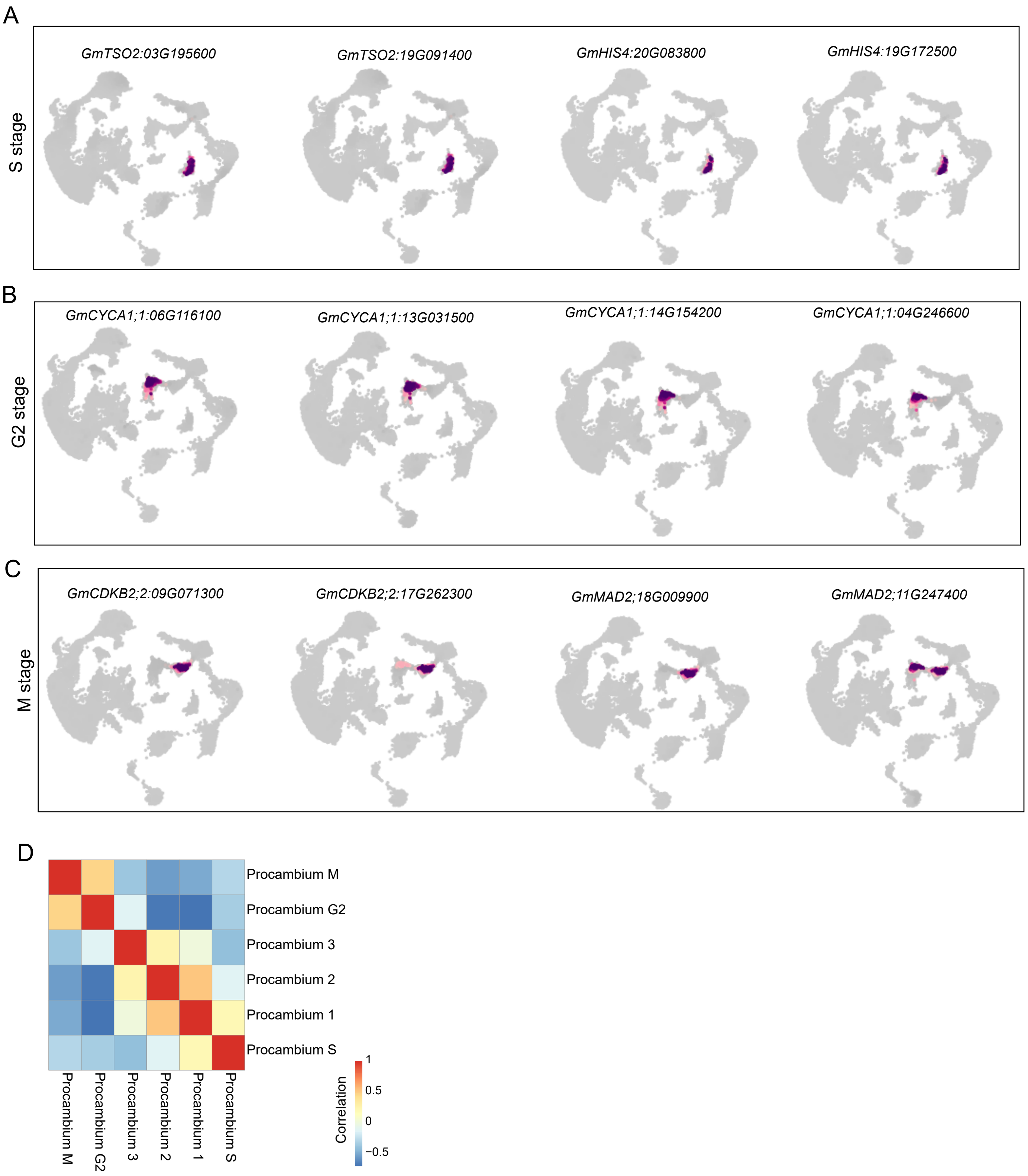


**Fig. S2 Cell cycle annotation of procambium cells**

(A-C) UMAPs showing expression of representative markers for S phase (A), G2 phase (B), and M phase (C).

(D) Heatmap showing Spearman's correlation coefficients between the 1,000 most variable genes from procambium cell types.
