## Supplementary material for "Single-cell multi-omic characterization of the soybean root response to cyst nematode infection": Figure S3

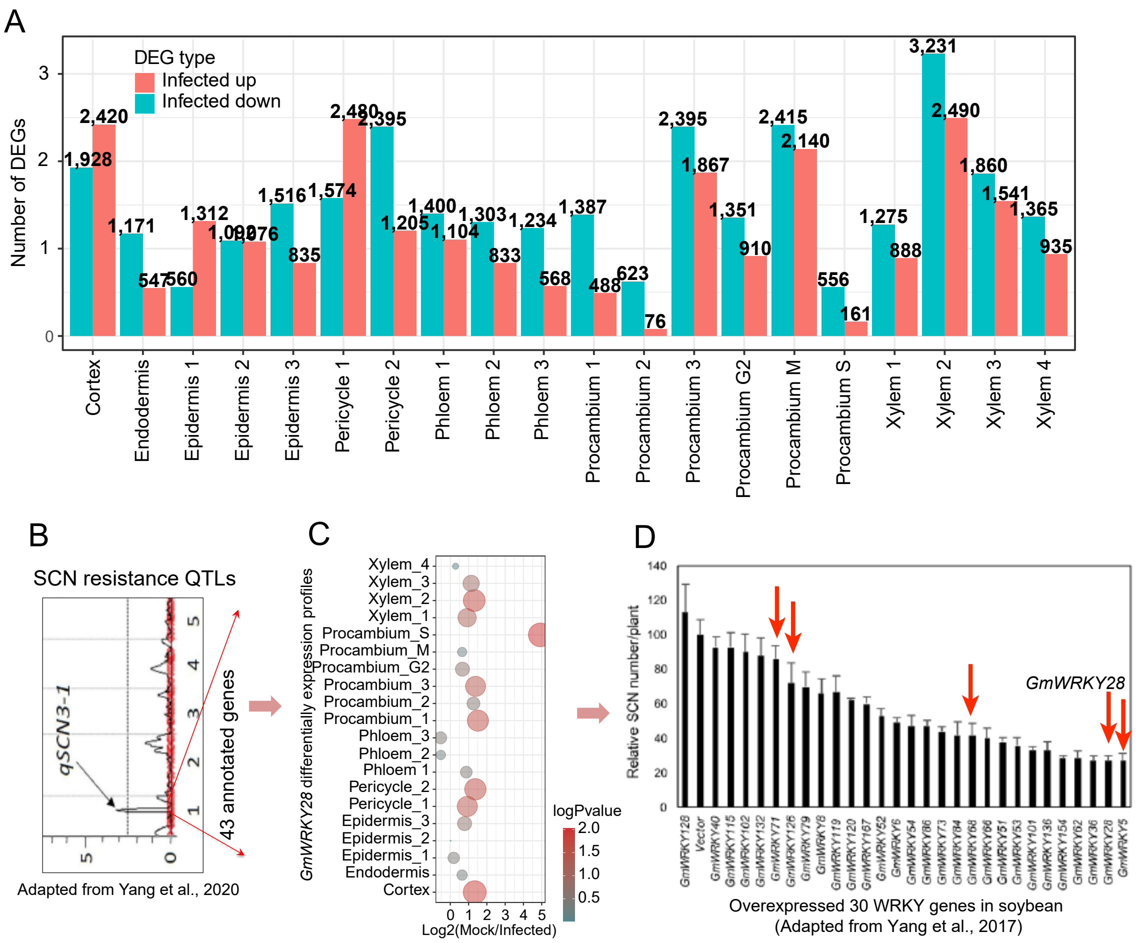

**Fig. S3 Identification and functional analysis of DEGs**

(A) Distribution of DEGs across all cell types between mock and infected roots.

(B) QTL mapping for SCN resistance (adapted from Yang *et al*., 2020)..

(C) Identification of *GmWRKY28* as a candidate gene for *qSCN3-1*, based on DEGs specific to Procambium S.

(D) SCN resistance assays in transgenic hairy roots overexpressing soybean *WRKY* genes (adapted from Yang et al., 2017). The five *WRKY* genes identified as DEGs in syncytium-associated procambium cells are marked with red arrows.
