## Supplementary material for "Single-cell multi-omic characterization of the soybean root response to cyst nematode infection": Figure S4

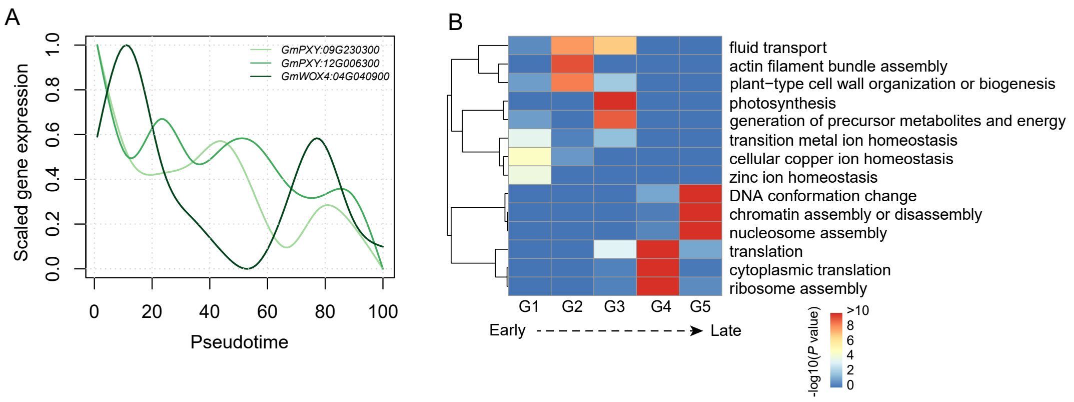


**Fig. S4 Functional characterization of pseudotime-associated genes**

(A) Expression patterns of *GmPXY (Glyma.09G230300, Glyma.12G006300)* and *GmWOX4 (Glyma.04G040900)* along the pseudotime trajectory.

(B) Three representative enriched GO categories for each of the five gene clusters (G1–G5).
